# The haplolethality paradox of the *wupA* gene in *Drosophila*

**DOI:** 10.1101/2020.09.11.292748

**Authors:** Sergio Casas-Tintó, Alberto Ferrús

**Author notes:** Correspondence: Alberto Ferrús.

## Abstract

Haplolethals (HL) are regions of diploid genomes that in one dose are fatal for the organism. Their biological meaning is obscure, given that heterozygous loss-of-function mutations will result in dominant lethality and, consequently, should be under strong negative selection. In addition, their biological nature is under debate: a chromosomal structure that is essential for the expression of several genes (regulatory hypothesis), or the strict dosage requirement of one particular gene (functional hypothesis). We report the first in depth study of an haplolethal region, the one associated to the *Drosophila* gene *wings up A* (*wupA*). It encodes 13 transcripts (A-M) that yield 11 protein isoforms (A-K) of Troponin I (TnI). They are functionally diverse in their control of muscle contraction, cell polarity and cell proliferation. Isoform K can transfer to the nucleus where it increases the transcription of the cell proliferation related genes *CDK2, CDK4, Rap* and *Rab5*. The nuclear translocation of isoform K is prevented by the co-expression of A or B isoforms, which illustrates isoform interactions. The corresponding dominant lethal mutations (DL) result from DNA rearrangements, clustered in the intron between exons 7-8, and affect the genomic organization of all transcripts. The joint elimination of isoforms C, F, G and H, however, do not cause DL phenotypes. Genetically driven expression of single isoforms rescue neither DL nor any of the mutants known in the gene, suggesting that normal function of *wupA* requires the properly regulated expression of specific combinations, rather than single, TnI isoforms along development. We conclude that the HL function at *wupA* results from the combined haploinsufficiency of a large set of TnI isoforms. The qualitative and quantitative normal expression of which, requires the chromosomal integrity of the *wupA* genomic region. Since all fly TnI isoforms are encoded in the same gene, its HL condition becomes unavoidable.

**Author summary:** Most species contain two copies of their genetic endowment, each received from their progenitors. If one of the duplicated genes is non-functional, due to either null mutations or genomic loss, the remaining gene copy may supply enough product as to cover the requirements for normal function or, alternatively, may reflect the insufficiency through a visible phenotype. In rare occasions, however, mutation in one copy is so deleterious that causes lethality, usually at early stages of development. These so called “haplolethal regions”, exist across species and represent an evolutionary paradox since they should have been subject to intense negative selection. The inherent difficulties to study haplolethals have precluded their study so far. Here, we analyzed the case of one of the five haplolethal regions of *Drosophila*, the one associated to the Troponin I encoding gene *wupA*, by measuring the transcriptional effects of mutations and chromosomal rearrangements affecting this gene. The data show that haplolethality results from the combined insufficiency of a large number of Troponin I isoforms, which are functionally specialized, show interference and require the integrity of the native chromatin structure for their quantitatively regulated expression. These features unveil novel aspects of gene expression and, possibly, on evolutionary gene splitting.

## INTRODUCTION

Diploid organisms are endowed with two genomic copies inherited from the parental generation. The gain or loss of small fragments from one or both of these genomic copies is referred to as segmental aneuploidies. When a single fragment is lost and the remaining copy is insufficient for normal biology, thus causing a phenotype, the segmental aneuploidy is referred to as haploinsufficient. In humans, haploinsufficient regions affecting to over 300 genes [1] are often associated to pathologies [e.g.: Parkinson’s disease [2]; bone marrow syndromes or myeloid malignancies [3] and autoimmune disorders [4], among others]. More rarely, haploinsufficiency may result in physiological advantages. In *S. cerevisiae*, a segmental haploidy increases stress resistance and ethanol production [5]. Interestingly, 3% of its 5900 genes are haploinsufficient when growing in enriched medium. The effect, however, can be alleviated by growing in minimal medium which suggests that the haploinsufficiency is due to low protein production [6]. Noticeably, 120 genes have never been recovered as deletions, suggesting that the absence of one copy of these genes may be lethal in the diploid phase of yeast [7]. Isolating such deletions is only feasible in diploid organisms by using suitable deficiencies and duplications. These regions are properly named as haplolethals (HL). Contrary to haploinsufficient regions, HL is a class of segmental aneuploidy whose nature remains unexplored due to the inherent difficulties to study them.

To justify a study on HL functions one should question how general HLs are? The seminal work by the groups of Lindsley and Sandler in 1972 [8], based on segmental aneuploidies covering 85% of the *Drosophila* genome, provided the first estimation of HL regions, up to 20, one of them being haplo-, as well as, triplo-lethal (Tpl). Since organism survival is inversely proportional to the extent of the genetic material deleted, the HL condition could result from the additive insufficiency of several adjacent genes. Thus, the estimated number of HL regions in flies has been reduced when smaller deletions have become available. A recent study of 793 small deletions covering 98.4% of the *Drosophila* genome indicates the number of HL regions as 5, including the Tpl [9]. The difficulty to study HL functions has precluded linking HL regions to specific genes, with only two exceptions. The *decapentaplegic* (*dpp*) gene includes an HL region which was functionally dissected from a recessive lethal phenotype using specific DNA fragments as transposons [10]. The HL region of *dpp* is referred to as Hin (haploinsufficient) in the corresponding literature. It spans 8 kb and it is thought to affect the five *dpp* transcripts. This Hin region seems refractory to insertions of the *P* type, but not to *hobo* type, transposons. Interestingly, the two 13 kb *hobo* inserts in the Hin region of *dpp* that had been reported, one of them landed in an intron, and the other did it in the 3’ untranslated region of exon 3 [11] (See **Suppl. Fig S1** which reproduces Fig 6 of [11]). Unfortunately, the HL component of *dpp* has remained uninvestigated, in comparison to the biology of the encoded Dpp/BMP protein, and the bases for the HL function at *dpp* are still unknown. The *wings-up A* (*wupA*) gene is related to another HL region located at chromosome band 16F7 [12]. An haplolethal region implies the existence of dominant lethal loss-of-function mutations. Intriguingly, the mutational analysis of the region searching for the expected dominant lethal mutants (DL), had provided chromosomal rearrangements only, rather than point mutations, and all of them turn out to be located towards the 3’ end of the *wupA* transcription unit [13]. Remarkably, the Tpl region has proven also refractory to point mutations and only rearrangements were obtained [14]. Since the duplication required to isolate DL mutants in the 16F7 region, *Dp*(*1;3R*)*JC153*, also contains genes adjacent to *wupA*, the HL function could result from the combined insufficiency of several genes under the control of regulatory sequences located inside the *wupA* gene or, alternatively, from the haploinsufficiency of the *wupA* encoded protein, Troponin I (TnI). Although we have previously analyzed the biology of TnI beyond the muscle cells [13,15–19], we address here the functional specificity of its various isoforms with the aim of unraveling the genetic nature of the associated HL.

In the mouse, one HL function was identified by inducing segmental haploidy in ES cells followed by their injection in blastocysts. It was ascribed to the *t*-complex gene [20–22]. The HL function was assigned to a 3Mb region which includes several genes and, consequently, the correspondence with one or several genes was not unequivocally established. However, two other murine HL functions, also generated by treated ES cells, could be associated to the *Vegf* [23] and *Tcof1* [24] genes. Based on theoretical considerations, HL functions in the mouse are estimated as more likely to occur than haploinsuficient ones (see Fig. 6 in [1]). In humans, about 1% of all pregnancies include some form of deviation from normal diploidy [25], and chromosomal deletions of embryonic malformations data revealed that about 11% of the genome has never been recovered in the haploid condition or any other copy number variation [26–29]. Although small deletions in human genomes are recognized as largely underestimated [30], segmental aneuploidies have been identified in oocytes, 10.4%, raising to 24.3% by three days of embryogenesis, and declining to 15.6% in preimplantation blastocysts [31]. Thus, human genomes reveal their instability, from meiosis to adulthood and, consequently, the need for continuous selection against aneuploidies. These observations could be taken as indirect evidence for deleterious HL functions in humans and its strong negative selection. In brief, the available data on HL functions, although scant, indicate that this type of mysterious functions are present across species, hence worth studying as a general feature of genomes.

Changes in gene dosage are sometimes compensated by transcriptional changes in functionally related genes outside the deleted region [32]. This effect is thought to reflect the robustness of the genome network [33]. In this context, HL functions could represent major failures in the chromatin topology required for proper transcription of several genes, either adjacent in a cluster or dispersed throughout the genome. We will refer to this interpretation as the *“regulatory hypothesis for HL”*. As an alternative, HL functions could correspond to single genes whose encoded product(s) are required in such a strict stoichiometry that a 50% dosage decrease becomes lethal. We will refer to this interpretation as the *“functional hypothesis for HL”*.

## RESULTS

This study deals mainly with the HL function located at the *Drosophila* chromosome band 16F7 where the gene *wupA* is located. Although the original data are available in previous reports [12,16,34], we summarize here the mutational analysis of *wupA* to facilitate reading this report and accommodate the novel data. Four categories of *wupA* mutations can be identified (**Fig. 1**). Described from proximal to distal with respect to the centromere, the mutations are classified as Recessive lethals (RL), Semidominant lethals (SDL), Dominant lethals (DL) and Viables (V). The types RL, SDL and DL are discriminated by the viability of heterozygous females, 100% in RL, under 50% in SDL and 0% in DL. While V mutations are single nucleotide changes, the RL, SDL and DL types are all chromosomal rearrangements. The transcriptional products from *wupA* result from two ATG translation sites (blue and red in **Fig. 2**) yielding 13 different transcripts, A-M, which are shown color-coded in **Fig. 2**.

**Fig 1.**
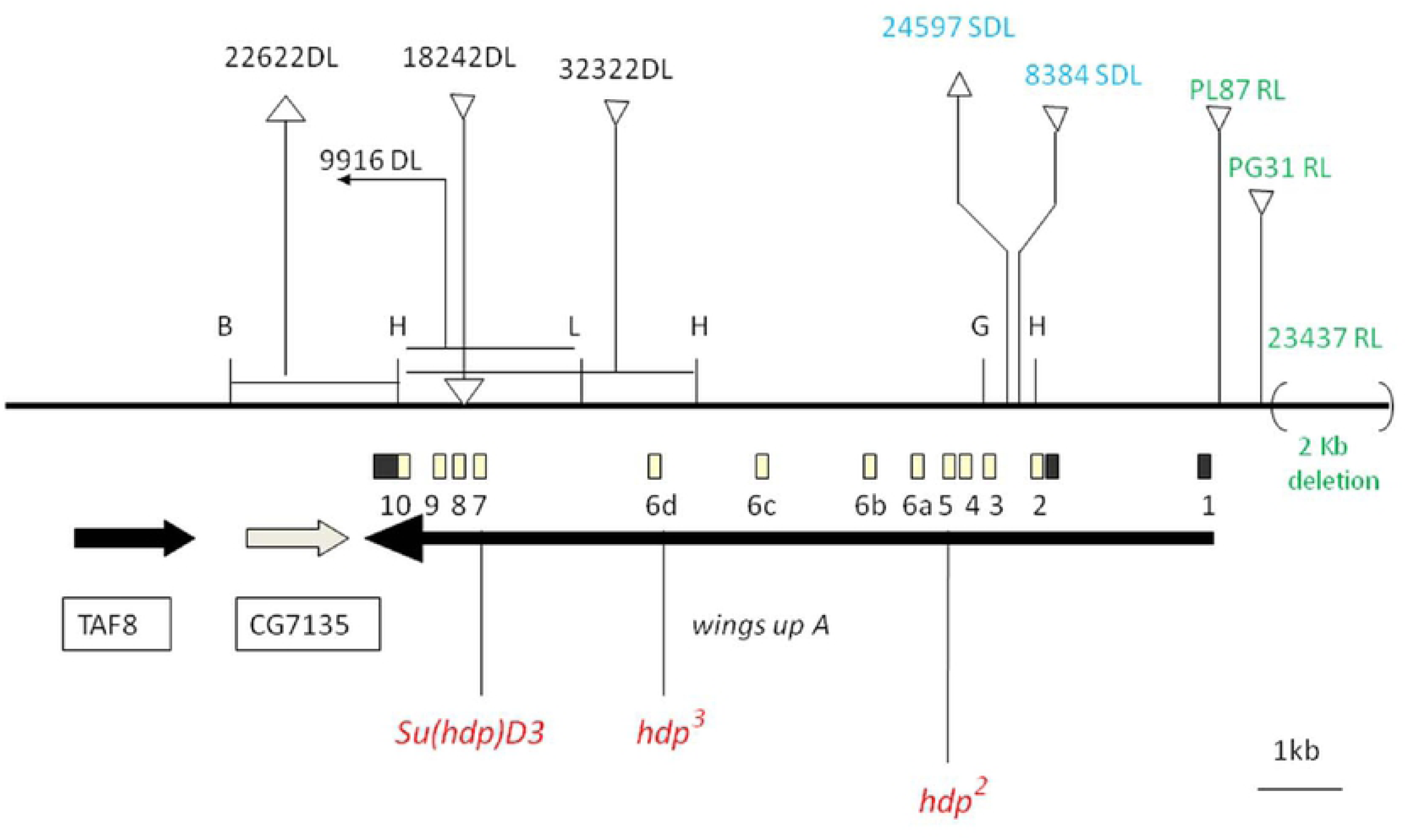
Mutations in the *wupA* gene. The diagram shows the relative positions of the *wupA* transcription unit (thick black arrow). Exons (1-10) are indicated by open (coding) and black (non-coding) boxes. Centromere is to the right. Transcription units *CG7135* and *TAF8* (a.k.a. *prodos*) locate distal to *wupA* and are indicate by thick arrows. Dominant lethal (DL) mutations are rearrangements whose breakpoint positions are determined by Southern blots with respect to enzyme restriction sites (B = *Bam*HI; H = *Hind*III; L = *Sal*I; G = *Bgl*II). *22622DL* is a deletion of the chromosomal segment 16F-18D. *32322DL* is an insertion of 23 Kb. *9916DL* is an inversion. The simplest of the DL mutations known to date, *18242DL*, is a 540 bp insertion in the intron between exons 7 and 8 (sequence deposited in EMBL X58188). Semidominant (SDL) rearrangements are indicated in blue. *24597SDL* is a deletion of about 0.4 Kb while *8384SDL* is an insertion of about 8 Kb. Recessive lethal (RL) rearrangements are shown in green. *23437RL* is a 2 Kb deletion located at position −100 upstream of the transcription initiation site in *wupA. PG31RL* is a 11.279bp insertion located at −249 with respect to the same initiation site. *PL87RL* is another 10.691bp insert at position - 30. All these RL rearrangements affect regulatory regions of *wupA* and severely reduce, but do not abolish, *wupA* transcription [17]. Point mutations are indicated in red. Mutant *hdp*^*2*^ is an 116Ala>Val substitution in the constitutive exon 5; *hdp*^*3*^ is a one nucleotide change affecting the acceptor 3’ splice site for exon 6d which prevents the expression of TnI isoforms C, F, G and H (see **Fig. 2**); and *Su*(*hdp*^*2*^)*D3* is an 188Leu>Phe substitution in the constitutive exon 7. Data are from Barbas *et al*.,1993; Prado *et al*., 1995; Prado *et al*., 1999.

**Fig 2.**
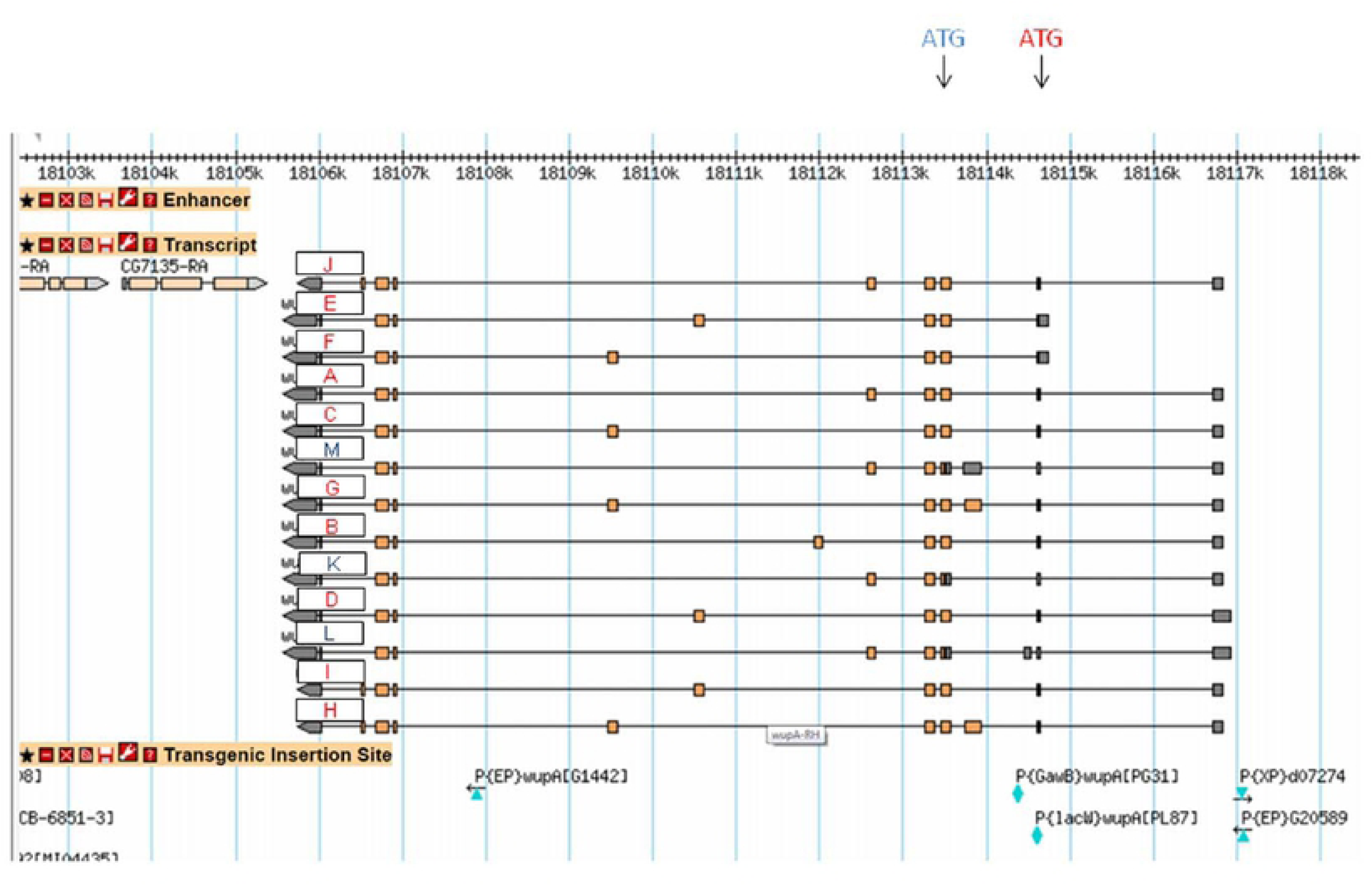
Transcriptional isoforms from *wupA*. The diagram shows the 13 transcripts emerging from the two ATG sites (red and blue). Note that the three blue transcripts (K, L and M) share the same protein coding exons, thus, the gene yields 11 TnI protein isoforms. Data are available in FlyBase.

### The normal HL function at 16F requires the integrity of the *wupA* genomic region

Over the years, a number of duplicated genomic fragments of the *wupA* region have become available (**Fig. 3**). In order to establish a link between the HL function and genes in the region, each duplicated fragment was tested against all available mutants in *wupA* and adjacent genes, in particular all known DL mutants. Green fragments indicate rescue of DL and *wupA* mutants, and red ones indicate fail to do so. Duplication 3 [Dp3 = *Dp*(*1,3R*) *JC153*] is the largest fragment (>600 kb) and, as expected, it fully rescues all DL and the rest of *wupA* mutants. Duplication 1 [Dp1 = *Dp*(*1;2R*) *CH322-143G12r*], a pBAC with which we obtained a transgenic line, also rescues all *wupA* mutants. Duplication 2 [Dp2 = *Dp*(*1;3*) *wupA-2XTY1-sGFP-V5-preTEV-BLRP-3XFLAG*] is a fosmid construct [35]. Particularly relevant is the result that the combination of genomic fragments *E6L* plus *Dp* (*1;2L*) *CH322-61E02r* is still unable to rescue DL mutants even though it encompasses the entire *wupA* transcription unit, (albeit not in a continuous sequence) (**Fig. 3**). If a regulatory sequence would be contained within the *wupA* transcription unit, it would not be able to operate in *trans*. Thus, the normal HL function at 16F7 requires the linear (*cis*) integrity of the *wupA* gene. In addition, the fact that ♀ DL/RL; Dp2/+ is lethal, but ♀DL/RL; Dp2/Dp2 is not, demonstrates that the RL and DL functions are functionally related and dosage dependent on the products supplied by Dp2 (see foot notes in **Table 1**).

**Table 1.** GENOTYPES VIABILITY

| **Genotype** | **Viability** |
| --- | --- |
| ♂ DL; Dp1/+ | YES |
| ♀ DL/+ ; Dp1/+ | YES |
| ♀ DL/DL ; Dp1/+ | NO |
| ♀ DL/DL ; Dp1/Dp1 | <1% |
| ♂ RL ; Dp1/+ | YES |
| ♀ DL/RL ; Dp1/+ | YES |
| ♂ hdp ; Dp1/+ | + |
| ♂ DL; Dp2/+ | NO |
| ♂ DL; Dp2/Dp2 | YES |
| ♀ DL/+ ; Dp2/+ | YES |
| ♀ DL/DL ; Dp2/+ | NO |
| ♀ DL/DL ; Dp2/Dp2 | <1% |
| ♂ RL ; Dp2/+ | YES |
| ♀ RL/RL ; Dp2/+ | YES |
| ♀ DL/RL ; Dp2/+ | NO |
| ♀ DL/RL ; Dp2/Dp2 | YES |
| ♂ hdp ; Dp2/+ | hdp |

Notes.- Viability determined from crosses in which a minimum of 100 progeny were counted (see M&M). Genotypes yielding escapers (<1% viability) consisted in short lived adults with normal gross morphology but unable to move properly or fully expand their wings and legs.
Abbreviations:
DL (dominant lethal) = *l(1)18242<sup>DL</sup>* or *l(1)13193<sup>DL</sup>*
RL (recessive lethal) = *l(1)23437*
hdp = *hdp<sup>2</sup>* or *hdp<sup>3</sup>*
Dp1 (duplication type 1) = *Dp(1;2R) CH322-143G12r*
Dp2 (duplication type 2) = fosmid, *wupA-2XTY1-sGFP-V5-preTEV-BLRP-3XFLAG*

**Fig 3.**
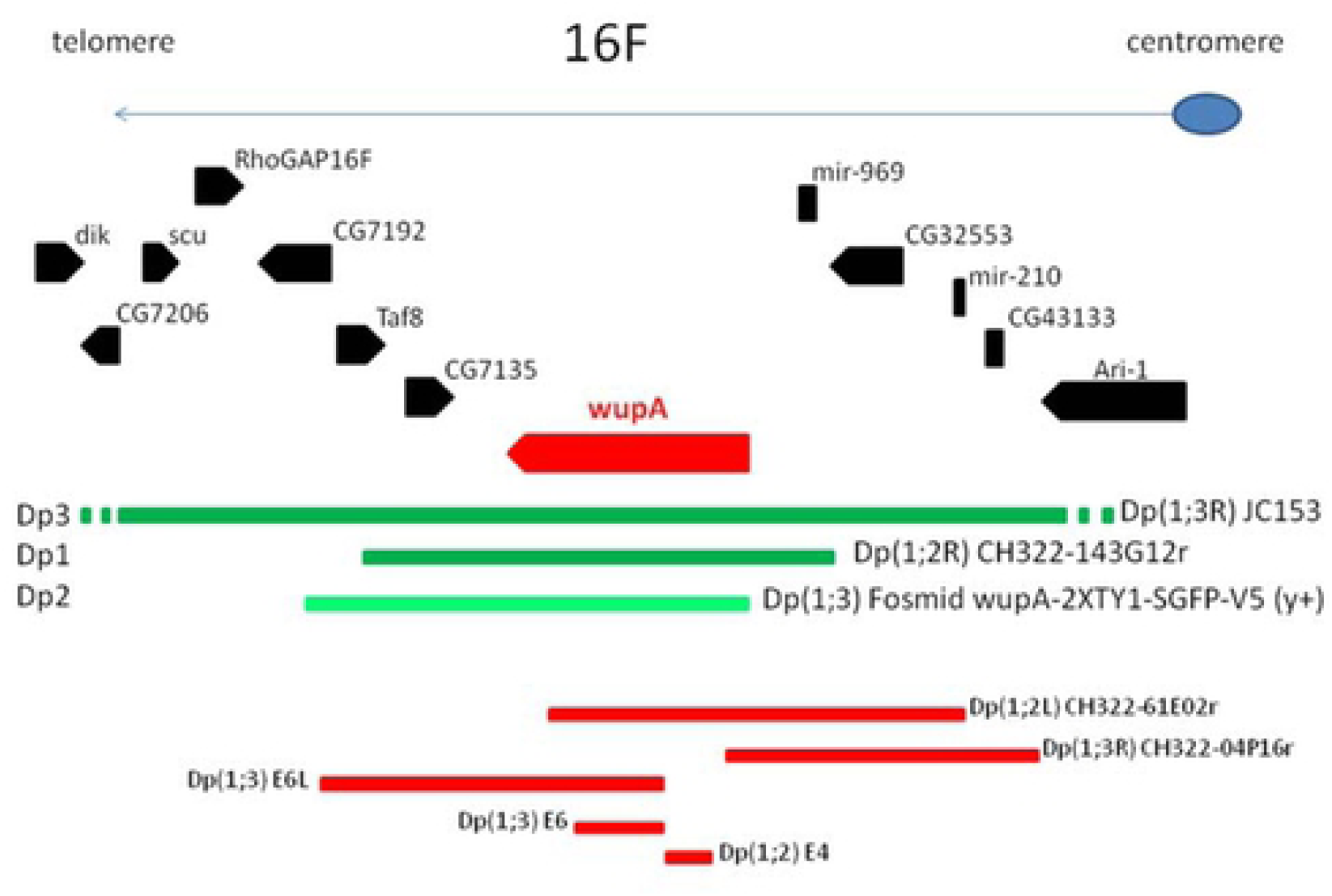
Duplicated chromosomal fragments covering the *wupA* region. Known transcription units distal and proximal to *wupA* are shown as pointed boxes. Thick lines illustrate the functional extent of the various duplications tested. Those in green cover all dominant lethal mutations while those in red do not. Duplications *E6L, E6* and *E4* correspond to genomic fragments inserted in the corresponding chromosomes. *E6L* spans 8 Kb and covers mutations in *Taf8* while E6 and E4 do not rescue any of the available mutants in the region. Duplications CH322 correspond to three pBAC constructs from which we obtained transgenic lines (see M&M). Only *CH322-143G12r* rescues DL mutants. This duplication is abbreviated as Dp1 in **Table 2**. The fosmid duplication *Dp*(*1;3*) *wupA-2XTY1-SGFP-V5* [35,66] covers mutations in *Taf8* and, most but not all, mutations in *wupA* (see main text). This fosmid duplication is abbreviated as Dp2 in **Table 2**. Duplication JC153 spans 550 Kb and covers all known mutants of the region. It originates from the insertional translocation (*T1;3*) *JC153*. This duplication is abbreviated as Dp3 in **Table 2**.

The adjacent location of *mir-969* (http://flybase.org/cgi-bin/gbrowse2/dmel/?Search=1;name=FBgn0283471) to *wupA* and its inclusion in Dp1, a duplication that covers the DL mutants (**Fig. 3** and **Table 1**), invited to explore its potential role in our HL function. Thus, *mir969* was expressed under the general driver *tub-Gal4*^*LL7*^ in normal, RL *l*(*1*)*23437* and V *hdp*^*3*^ backgrounds. In the first case, the overexpression of *mir969* throughout the body did not yield a visible phenotype. In the other two backgrounds, it failed to rescue or modify, either of these *wupA* mutations. Likewise, the depletion of *mir969* by means of expressing the construct *UAS-mCherry-mir969-sponge* under either of these three backgrounds, also failed to modify the *l*(*1*)*23437*^*RL*^ and *hdp*^*3*^ phenotypes nor to cause a DL condition. Most relevant, it should be noted that Dp2, which rescues the DL mutants, does not contain *mir-969* (**Fig. 3**). Thus, we can conclude that *mir969* does not show evidences to suspect a functional interaction with *wupA* or its associated HL function. An equivalent reasoning allows excluding the other adjacent gene, *CG7135*, since the fragment E6L does not rescue any of the *wupA* mutants (**Fig. 3**).

**Table 2.** RESCUE OF wupA MUTANTS

|  | DL |  | SDL | RL |  |  | V |  |
| --- | --- | --- | --- | --- | --- | --- | --- | --- |
|  | 18242<sup>DL</sup> | 13193<sup>DL</sup> | 24597<sup>SDL</sup> | 23437<sup>RL</sup> | PL87<sup>RL</sup> | PG31<sup>RL</sup> | hdp<sup>2</sup> | hdp<sup>3</sup> |
| **A** | - | - | - | - | - | - | - | - |
| **B** | - | - | - | - | - | - | - | +<sup>(b)</sup> |
| **C** | - | - | - | - | - | - | - | - |
| **D** | - | - | - | - | - | - | - | - |
| **E** | - | - | - | - | - | - | - | +<sup>(b)</sup> |
| **F** | - | - | - | - | - | - | - | - |
| **G** | - | - | - | - | - | - | - | - |
| **H** | - | - | - | - | - | - | - | - |
| **I** | - | - | - | - | - | - | - | - |
| **J** | - | - | - | - | - | - | - | - |
| **K** | - | - | - | - | - | - | - | - |
| Dp1 | + | + | + | + | + | + | + | + |
| Dp2 | +<sup>(a)</sup> | +<sup>(a)</sup> | + | + | + | + | - | - |
| Dp3 | + | + | + | + | + | + | + | + |

**Notes.-** The wupA transcripts are depicted in red or blue (A-K) according to their corresponding ATG site (see **Fig. 2**). Every isoform was driven as UAS constructs by tub-Gal4 and a minimum of 100 offspring adults per cross were counted. Blue transcript isoforms K, L and M encode the same protein, thus only K was assayed. (a) Rescue is dose dependent; one copy of Dp2 can rescue ♀DL/+ but two copies are needed to rescue ♂DL or ♀DL/DL. In the last case, only escapers are recovered as adults (<1% viability). The same driver was used to co-overexpress several transcripts (C+H, J+F and J+G) but none of these combinations rescued 18242<sup>DL</sup> neither in males nor in heterozygous females. (b) Rescue of hdp<sup>3</sup> by either of these two transcripts was obtained for 25% (B) and 20% (E), respectively. The rescue, however, affected to wing position but not to flight. The joint overexpression of C and H also failed to rescue hdp<sup>3</sup>.
Dp1 (duplication 1) = Dp(1;2R) CH322-143G12r
Dp2 (duplication 2) = fosmid, wupA-2XTY1-sGFP-V5-preTEV-BLRP-3XFLAG
Dp3 (duplication 3) = Dp(1;3R)JC153

In addition to the HL functions linked to *wupA* and *dpp* genes, there are other genomic regions for which the deletion analysis indicates that HL functions may exist. We explored a possible interaction between the *wupA* HL and the rest of putative HL functions, excluding the Tpl region due to the lack of suitable genetic tools (**Suppl. Table S1**). We tested genotypes carrying a DL mutant at *wupA* and duplications that cover the other reported HL regions [36,37]. Likewise, we tested deficiencies that uncover these other HL regions with duplications that cover *wupA*. None of these combinations modified the corresponding HL phenotypes. Thus, we conclude that there is no evidence, at this point, to suggest a functional link among HL functions within the *Drosophila* genome. They do share, however, the coding of multiple transcripts and, in the known cases of *dpp* and *wupA*, the corresponding DL mutants are rearrangements broken in non-coding sequences.

Taken together, this set of data strongly support the hypothesis that the HL function at chromosome band 16F7 results from the haploinsufficiency of products encoded in *wupA* exclusively. Nevertheless, why all known DL mutations are chromosomal rearrangements?

### Single transcripts from *wupA* rescue neither DL nor *wupA* mutants

UAS constructs were generated for each *wupA* transcript and tested with the general driver *tub-Gal4*^*LL7*^ in order to assay if the overexpression of any of them would rescue mutants located in this gene. The data show (**Table 2**) that none of the red transcripts, nor the single blue one tested, rescued any of the *wupA* mutant types including the DL type. The rescue of *hdp*^*3*^ by isoforms B and E is partial since they restore wing position, albeit not flight, in about 25% of adults only. At least two different transgenic insert lines were tested per UAS construct.

We also tested three relevant genomic duplications with the same set of *wupA* mutants in order to compare with the results obtained with the UAS constructs (**Table 2**). Duplication 2 yielded interesting rescue effects. It shows dose dependence in the rescue of DL mutants. One copy of Dp2 is sufficient to rescue DL/+ heterozygous females, while two copies are needed to barely rescue DL/DL homozygous adult females. Two copies are also needed to rescue DL males. Another interesting feature of Dp2 is its failure to rescue the V type mutants *hdp*^*3*^ and *hdp*^*2*^. This last result demonstrates that Dp2, although able to rescue DL mutants, does not supply the full set of normal functions for *wupA*.

The two sets of data, the lack of rescue by the single *wupA* transcripts and the efficient rescue by some genomic duplications, could be taken as evidence in favor of the regulatory hypothesis; that is, a putative regulatory structure operating in *cis* to control the proper expression of the TnI isoforms. Two of the duplications, however, showed differential rescue effects, Dp1 and Dp2. This feature invited to analyze the transcriptional properties of these two duplications. To that end, we used qRT-PCR to measure the transcriptional yield of adult males with Dp1 and Dp2 in *l*(*1*)*23437* and *hdp*^*3*^ backgrounds. These backgrounds and the designed exon specific probes facilitated the identification of *wupA* transcripts to the extent possible (**Fig. 4**). In the *l*(*1*)*23437* background (**Fig. 4A-D**), the heterogeneity in the relative levels of the various transcripts, either in the presence of one or two copies of Dp2 or in combination with Dp1, becomes evident. In the *hdp*^*3*^ background (**Fig. 4E-G**), the phenomenon of quantitative transcriptional heterogeneity is also detected. Even the point mutation *hdp*^*3*^ that, altering the splice site for exon 6d should have affected the C, F, G and H red transcripts only, yields overexpression of red A, J, and blue K, L and M products (**Fig. 4E**). Neither Dp1 nor Dp2 can normalize this *hdp*^*3*^ caused overexpression. These transcriptional data uncover a fine quantitative regulation of the *wupA* transcriptional expression that was unsuspected hitherto. Not only the genomic duplications do not provide the expected levels of transcripts, but also a splice site mutation as *hdp*^*3*^ seems to alter the expression of transcripts that do not include the affected exon, 6d. In addition, these data also suggest that the *wupA* encoded products, isoforms of Troponin I (TnI), may not be functionally equivalent.

**Fig 4.**
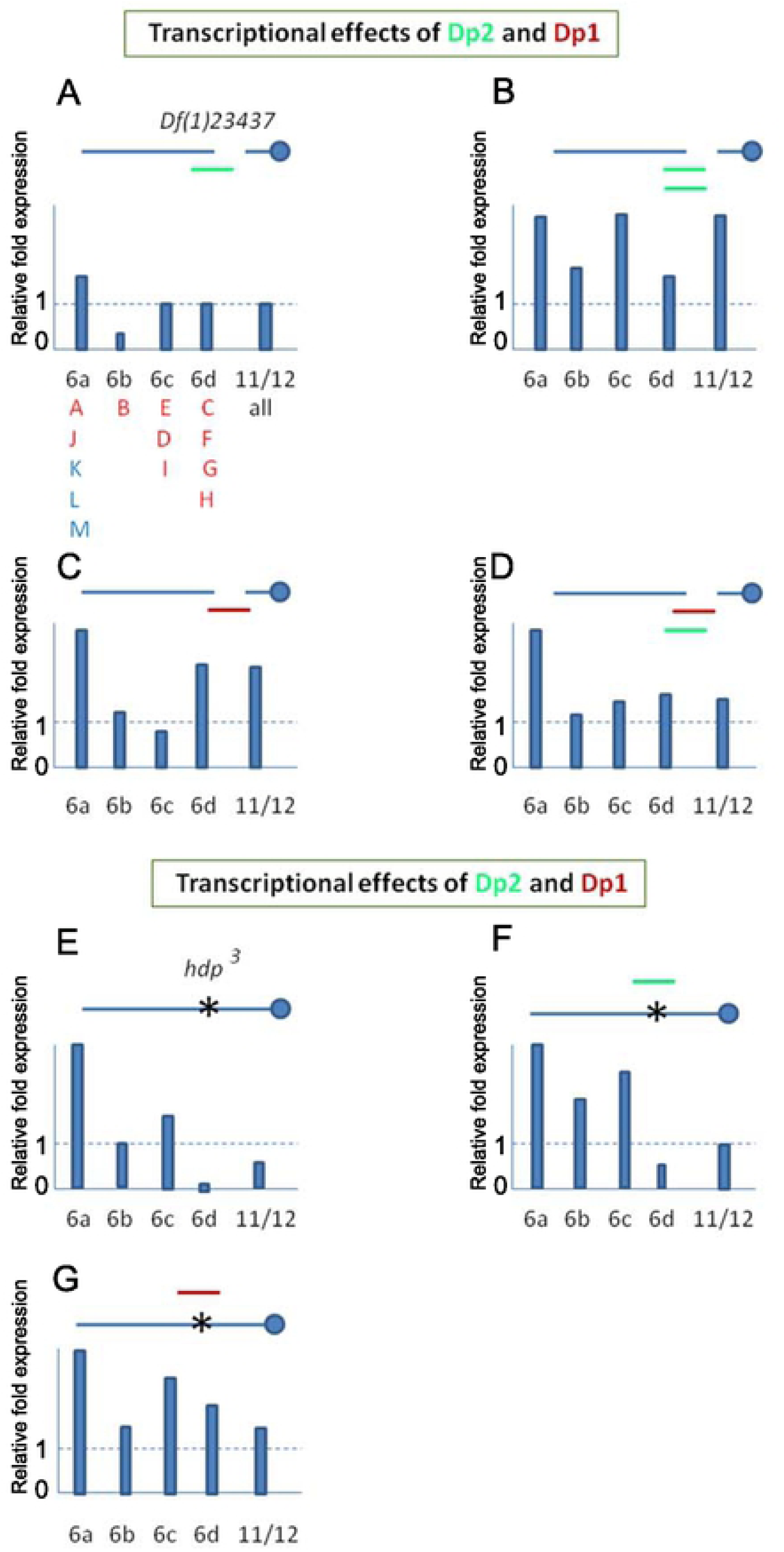
Transcriptional effects of two duplications covering *wupA*, Dp1 and Dp2. A-D) Effects on *l*(*1*)*23437* background. We tested the transcriptional efficiency of these two duplications using exon 6 specific primers in qRT-PCR assays on severe hypomorph (*23437*) males. The *wupA* transcripts identified by each exon-specific primer are indicated in **A**. Dotted line indicates the normalized levels of controls. Note that one dose of *fosmid* (Dp2) fails to produce normal levels of isoform B and two doses do not duplicate linearly the transcription of all isoform. A similar transcriptional heterogeneity is observed with *Dp*(*1;2R*)*CH322-143G12r* (Dp1) in one dose or in combination with Dp2. **E-G) Effects on *hdp***^***3***^ **background**. The same heterogeneity is detected on this background. Note, however, that, in general, the transcriptional efficiency of Dp1 is higher than that of Dp2. This is consistent with the rescue data shown in **Table 2**.

### The *wupA* products are functionally diverse and interacting

In a previous study we had shown that the attenuation of *wupA* expression by means of RNAi or its overexpression using the TnI overexpressing construct *PBac*(*WH*)*f06492* yield noticeable effects on cell proliferation [19]. This overexpression is synergistic with standard oncogenes such as *Ras*^*v12*^, *Notch* or *lgl*, while its attenuation largely suppresses their tumor overgrowths. Following the generation of UAS constructs for single *wupA* transcripts, we analyzed the effects of their overexpression in order to determine if all TnI isoforms play similar functions in cell proliferation or if, by contrast, they show functional specificities.

To that end, we generated somatic clones in larval wing discs and quantitated clone size with respect to wing disks of sibling controls expressing an innocuous UAS construct (**Fig. 5A**). The data reveal at least three different phenotypic effects on cell proliferation: 1) No effect (transcript A), 2) Underproliferation to various degrees (transcripts B-J), and 3) Overproliferation (transcript K). Thus, different TnI isoforms affect cell proliferation and/or survival in different ways. While most isoforms reduce cell proliferation, isoform K causes a strong proliferative effect when overexpressed. Of notice, the three blue transcripts K, L and M encode the same protein which justifies testing only one of them.

**Fig 5.**
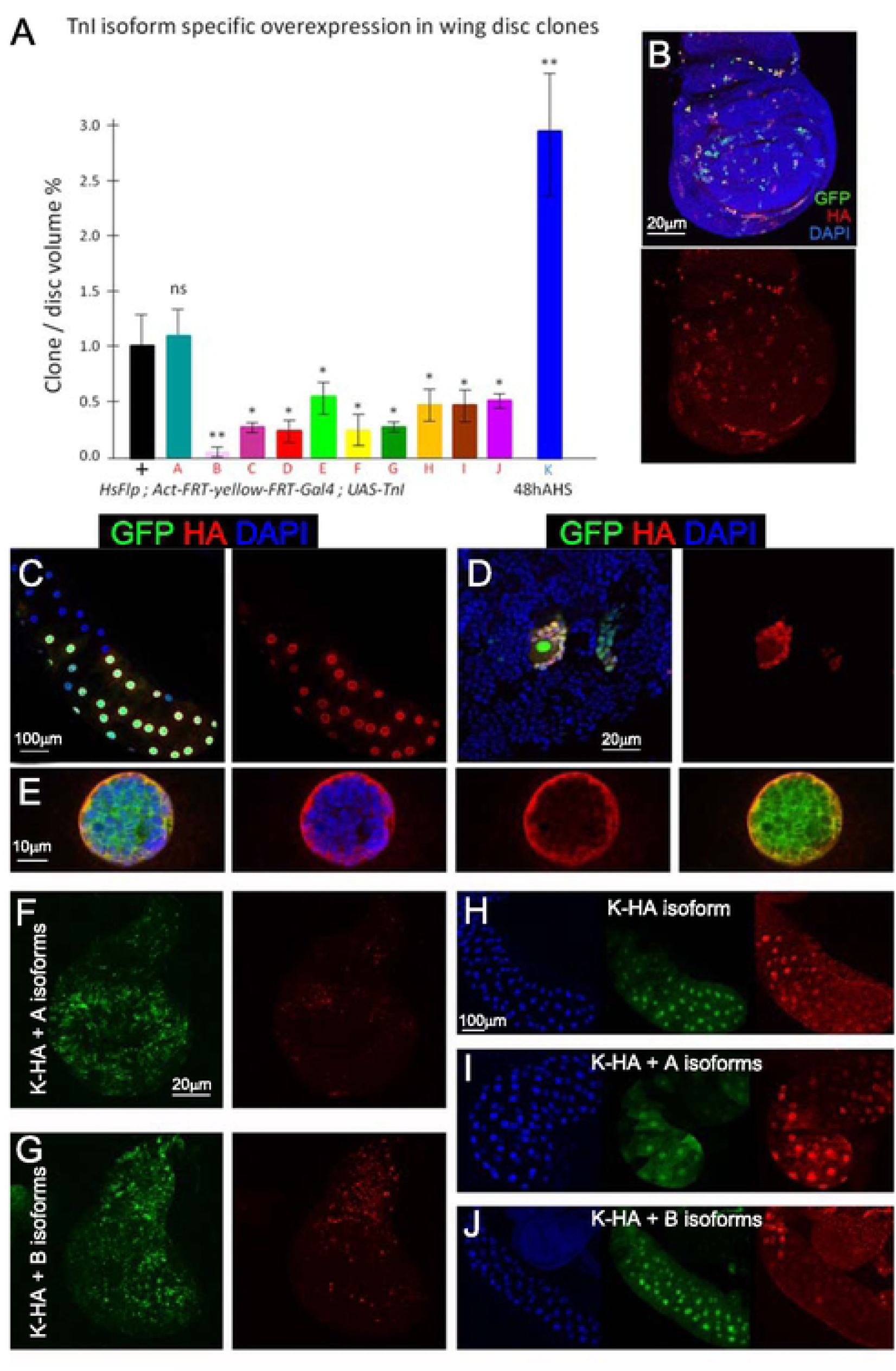
TnI isoforms are functionally different and interacting. A) Cell proliferation effects of single TnI isoforms overexpression. Somatic clones were induced in the larval wing discs (genotype: ♀♂ *hsFlp*; *act-FRT-yellow-FRT-Gal4/*+; *UAS-TnI*^*isoformX*^/+) screened 48h after heat shock (48AHS). Note that, with the exception of A, all red isoforms reduce cell proliferation. By contrast, the blue isoform tested (K), yields a significant proliferation increase. Since all blue isoforms (K, L and M) yield the same protein, only one was tested. **B-E) Nuclear localization of isoform K. B)** Images of larval wing disc somatic clones expressing the HA-tagged K isoform (red) and the cell reporter GFP to identify the clones. Nuclei are stained by DAPI (blue). **C)** Equivalent clones in the salivary glands and **D**) the neuroblast descendants in the central nervous system. **E**) Lower row of panels show one salivary gland nucleus stained for DAPI (blue) and TnI (red). Note the nuclear localization of K-HA. **F-J) The nuclear localization of isoform K is interfered by isoforms A or B**. Equivalent clones co-expressing TnI isoforms K-HA and A or B in wing imaginal discs (**F, G**) or salivary glands (**H-J**). The nuclear localization of isoform K is efficiently prevented by isoform A and, to a lesser extent, by isoform B.

Also in a previous study, we showed by immunodetection that TnI traffics to the cell nucleus as a function of cell cycle status [18]. Since isoform K causes overproliferation and we had shown previously that the generalized overexpression of *wupA* triggers overexpression of cell division related genes [19], we investigated if isoform K, by itself, could have the capacity to translocate to the nucleus and trigger transcriptional changes in cell proliferation related genes. To that end, we created an HA-tagged version of isoform K and expressed it in wing disc somatic clones, salivary glands and neuroblasts. The HA tag is detected in the nucleus in all three cell types (**Fig. 5B-E**). Benefiting from the large size of salivary gland nuclei, the HA-K signal can be clearly identified in the periphery of the nucleus (**Fig. 5E**).

In addition, we tested the eventual changes in the expression of a set of genes involved in the control of cell proliferation by qRT-PCR (**Suppl. Fig. S2**). We used RNA from larvae overexpressing isoform K (genotype: *tub-Gal4*^*LL7*^ > *UAS-K*). Consistent with our previous study, a subset of these genes are overexpressed when the isoform K is in excess. In particular, *CDK2, CDK4, Rap* and *Rab5* exhibited significant overexpression with respect to controls (genotype: *tub-Gal4*^*LL7*^ > *UAS-LacZ*). *CD2* and *CDK4* are well known inducers of cell cycle entry [38]. For example, *CDK2* mediates *Myc* induced cell proliferation through its association with Cyclin E [39,40], while *CDK4* plays equivalent roles through its association with Cyclin D [41,42]. Likewise, *Rap*, a Fizzy-related protein, regulates cell proliferation [43], and *Rab5* contributes to proliferative cell signaling through the titration of EGFR [44], among other functions.

Given the strong effect of isoform K on cell proliferation (**Fig. 5A**) and its nuclear localization, we questioned if its generalized overexpression could yield a visible phenotype. Two constructs were tested, HA-K and non-tagged K, driven by *tub-Gal4*^*LL7*^. In both cases, the genotype was adult lethal. Other drivers with more restricted domains of expression, *en-Gal4* and *rn-Gal4*, yielded poorly viable adults (<10%) with various morphological abnormalities in wings and legs (**Suppl. Fig. S3**). Thus, we conclude that the permanent and generalized overexpression of isoform K is organism lethal.

In view of the diversity of TnI effects on cell proliferation (**Fig. 5A**), we questioned if the function of a given transcript could be modified by others from the same gene. As an example, we chose the nuclear localization of the K isoform (blue) under the co-expression of red isoforms. Thus, we generated somatic clones expressing HA-K along with A or B isoforms. The data show that both red isoforms prevent the nuclear localization of K in imaginal discs and salivary glands, being the A effect stronger than that of B (**Fig. 5F-J**). To further investigate possible additional examples of isoform interference, we tested the adult lethality caused by the excess of K (genotype: *tub-Gal4*^*LL7*^ > K), but generated from females heterozygous for the RL mutant *l*(*1*)*23437*. In this case, the overexpression of K is no longer lethal and females *l*(*1*)*23437/*+; *tub-Gal4*^*LL7*^*/K* are 100% viable. This result suggested that the relative depletion of TnI isoforms caused by the *23437* mutant in the maternal oogenesis or early embryogenesis could alleviate the deleterious overexpression of isoform K. The suppression of the lethality due to K excess is also observed when the maternal progenitor is heterozygous for other RL mutants, *PL87* and *PG31*. The V mutants *hdp*^*3*^ and *hdp*^*2*^ also suppress the K-dependent lethality, always when the RL or V mutants come from maternal, not paternal, origin. As shown above (**Fig. 4E**), *hdp*^*3*^ eliminates isoforms C, F, G and H, and *hdp*^*2*^ is a point mutation in a constitutive exon (**Fig. 1**). On the other hand, the co-overexpression of K with isoforms B or A maintains the lethality. Finally, as shown in **Table 2**, isoforms B or E can rescue, to some extent, the wings up phenotype of *hdp*^*3*^. That is, the depletion of C, F, G and H isoforms can be compensated, at least in part, by the excess of B or E (see foot note in **Table 2**).

Beyond these in vivo phenotypes, we investigated the transcriptional effects of overexpressing specific TnI isoforms. The rationale being that, if there is phenotypic interference among isoforms, the overexpression of one of them could alter the expression of others (**Fig. 6**). The data show that the overexpression of isoform A (**Fig. 6A**), in addition to yield its own excess, as revealed by probe 6a, it also causes excess of probe 6c detected isoforms, E, D and I. Surprisingly, driving isoform B (**Fig. 6B**), yields a significant excess of most other isoforms except of itself. This paradoxical result could indicate, perhaps, a regulatory effect of isoform B on *wupA* transcription but the issue was not investigated further. The equivalent experiment driving isoform K (**Fig. 6C**) seems to be rather specific for that isoform as revealed by probe 6a. The transcriptional effect was also reproduced by driving the HA-tagged version of isoform K (**Suppl. Fig. S4**). Finally, we included in this set of experiments the transcriptional effects of the *PBac*(*WH*)*f06492* since this UAS-dependent construct had been shown to cause strong proliferation effects synergic with standard oncogenes [18] as mentioned above. The data show (**Fig. 6D**) that this construct elicits the strong overexpression of 6a revealed isoforms, which include isoform K, and, to a lesser extent, the 6c revealed isoforms. This transcriptional effect is consistent with the cell proliferative phenotype observed with isoform K overexpression (**Fig. 5A**).

Taken together, these experiments reveal a wide range of functional interactions among TnI isoforms operating throughout development and cell types. Presumably, the repertoire of interactions will extend beyond the cases experimentally analyzed here. Their large combinatorial number, however, precludes an exhaustive analysis at this time. It seems that the normal biology of the *wupA* gene consists of a collection of diverse, albeit interacting, functions achieved by the ensemble of encoded TnI isoforms. Likely, the expression of these isoforms will be subject to a tight quantitative regulation among themselves. Could a combined depletion of all or several TnI isoforms account for the HL function at 16F?

### Targeting a subset of *wupA* products causes a DL phenotype

The functional diversity of TnI isoforms invited to consider their haploinsufficiency as the cause of the HL phenomenon. To address this possibility, we obtained a point mutation (A>C) at the red ATG site by means of the CRISPR/Cas9 system (see M&M and **Suppl. Fig. S5**). Two independent mutations, *18230C* and *18230B*, were isolated and tested for their viability effects in combinations with DL, RL and V mutants and the same three genomic duplications used in previous experiments. The key data for allele *18320C*, which was validated by sequencing, are shown in **Table 3** and **Fig. 7**.

**Table 3.** FUNCTIONAL PROPERTIES OF A Dominant Lethal MUTATION IN THE red ATG SITE

| Genotype | Viability |
| --- | --- |
| ♂ C ; Dp3/+ | YES |
| ♂ C ; Dp2/+ | YES |
| ♂ C ; Dp1/+ | YES |
| ♀ C/+ ; +/+ | NO |
| ♀ C/C ; Dp1/Dp1 | YES |
| ♀ C/C ; Dp1/+ | NO |
| ♀ C/DL ; Dp3/+ | <10% |
| ♀ C/DL ; Dp2/+ | NO |
| ♀ C/DL ; Dp1/+ | NO |
| ♀ C/RL ; Dp3/+ | YES |
| ♀ C/hdp<sup>3</sup> ; Dp2/Dp2 | hdp |

Notes.- Assays were carried out with two independent CRISPR/Cas9 mutations (ATG>XYZ) induced on the red ATG site (see **Suppl. Fig. S4**). Table shows the results for allele *18320C* which were identical to those from allele *18320B*. Allele *18320C* was validated by PCR and sequencing (see M&M and **Fig 6**). A minimum of 100 adult offspring per cross were counted.
Abbreviations:
C = (dominant lethal) *wupA*<sup>18320C</sup>
DL (dominant lethal) = *l(1)18242*<sup>DL</sup> or *l(1)13193*<sup>DL</sup>
RL (recessive lethal) = *l(1)23437*
hdp = *hdp*<sup>3</sup>
Dp1 (duplication type 1) = *Dp(1;2R) CH322-143G12r*
Dp2 (duplication type 2) = fosmid, *wupA-2XTY1-sGFP-V5-preTEV-BLRP-3XFLAG*
Dp3 = *Dp(1;3R)JC153*

**Fig 6.**
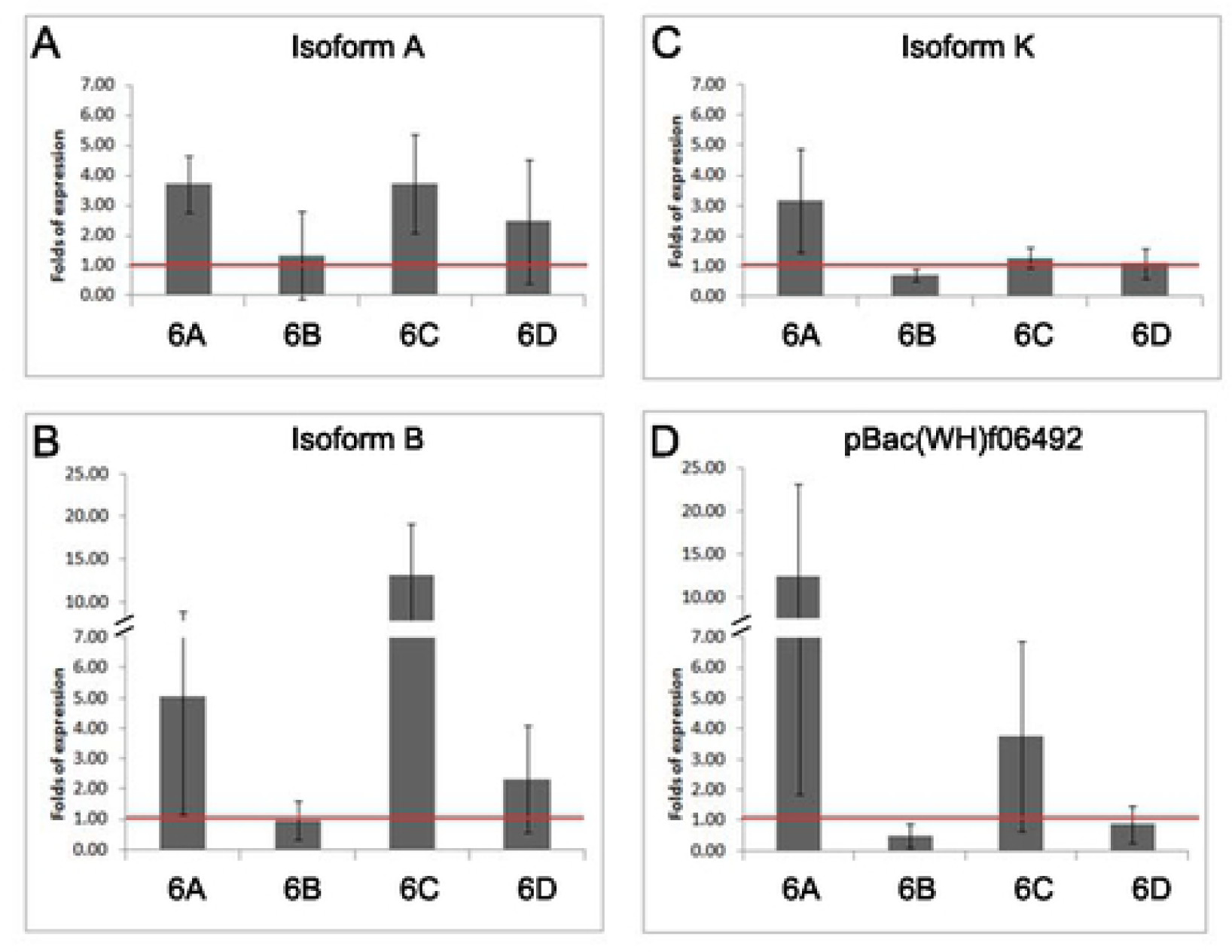
Transcriptional effect of driving single isoforms. **A-C)** Overexpression of isoforms A, B and K. **D)** Overexpression of the Exelxis construct for *wupA*. Note the systematic lack of effect upon the expression of isoform B which could suggest a regulatory role for this isoform (see main text). Genotype: ♂ *tub-Gal4*^*LL7*^>*UAS-TnI*^*isoformX*^. Primers from exons 6a-d identify TnI isoforms as indicated in **Fig. 4A**. Transcriptional levels are normalized to sibling controls ♂ *TM3/UAS-TnI*^*isoformX*^.

**Fig 7.**
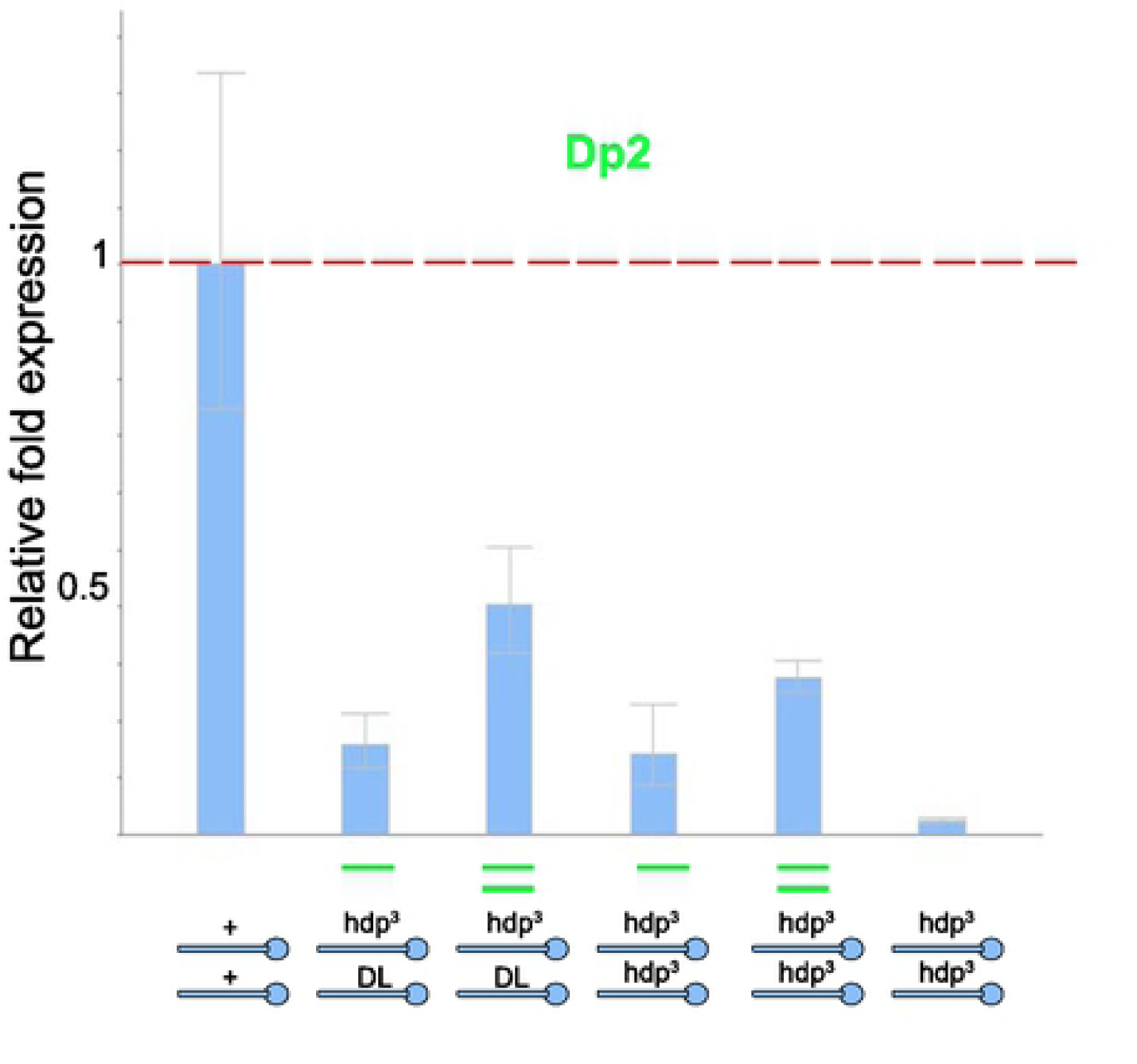
Transcriptional effects of the DL mutant allele *18320C*. Genotypes are indicated below each histogram. Primers from exon 6d were used to monitor isoforms C, F, G and H. Transcriptional levels are normalized to sibling controls. Note that *Dp2*, although it rescues the dominant lethality of *18320C*, it does not supply the normal levels of 6d containing isoforms, 25% roughly. Also, since *hdp*^*3*^*/hdp*^*3*^; *Dp2* and *hdp*^*3*^*/18320C; Dp2* yield the same levels of these transcripts, it follows that *18320C* is null for these isoforms.

Females heterozygous for *18230C* show 0% viability, thus proving its DL condition. This lethality is rescued by the three genomic duplications, Dp1, 2 and 3, when tested in males. In homozygous females, however, Dp1 shows the same dosage dependence that Dp2 had shown in previous experiments (**Table 2**). The *18230C* mutant, when confronted with regular DL mutants in heterozygous females, yields a lethality that neither Dp1 nor Dp2 can rescue. Actually, the large Dp3 yields a low rate of escapers only. By contrast, Dp3 does rescue *18230C* when heterozygous over a RL mutant. These data prove that the DL, RL and *18230C* mutants affect the same function(s) although, most likely, to a different degree. Also, the incomplete rescue by the duplications is consistent with their heterogeneous transcriptional yield shown above (**Fig. 4**).

To determine that *18230C* affects transcription of red isoforms, we carried out two experiments. On the one hand, adult females heterozygous for *18230C* and *hdp*^*3*^, whose dominant lethality is covered by Dp2, show the classical wings up phenotype (**Table 3**). On the other hand, the qRT-PCR assay of these flies (genotype: *18230C/hdp*^*3*^; *Dp2/*+) shows depletion of exon 6d containing isoforms, C, F, G and H; confirming that *18230C* eliminates these red isoforms (**Fig. 7**). Choosing Dp2 for these experiments is justified because it is the only available duplication that does not cover *hdp*^*3*^ and, yet, provides enough normal *wupA* function as to cover the dominant lethality. The quantitative transcriptional levels provided by Dp2 are consistent with the unusual dosage effects observed in the rescue experiments (**Table 2**). Either one or two copies of Dp2 fail to yield normal levels of the 6d reveled isoforms, C, F, G and H. Although it shows a clear dosage effect, two copies of Dp2 still transcribe below 50% of controls.

The data from the other allele, *18230B*, were identical to those of *18230C*. Targeting the blue ATG site would have not been instructive because, being located downstream of the red ATG, the procedure to obtain mutations by CRISPR in this triplet would also affect the red isoforms; thus providing no additional information with respect to the other existing DL rearrangement mutants.

These data demonstrate that the elimination of the whole red subset of TnI isoforms is sufficient to cause a DL phenotype; thus, it seems that the HL function at 16F7 results from the haploinsufficiency of TnI proteins, at least those encoded by the red transcripts. Considering all available *wupA* mutants and genomic duplications studied here, a graded array of product depletion becomes evident (**Fig. 8**). The most extreme is, obviously, the chromosomal deficiencies that uncover the region, and the DL rearrangements clustered at the 3’ end of the transcription unit. Following are the two *18230* mutants which also result in dominant lethality by eliminating the red isoforms only. The evidence that indicates that *18230* mutants are less severe than the regular DL mutants is shown in **Table 3**, where Dp3 can produce some escapers of the *18230C*/DL genotype. This Dp3, in one dose only, does not rescue DL/DL genotypes at all. Although not analyzed here, the next grade of depletion would be the SDL type of mutants, followed by the RL type. Finally, the *hdp*^*3*^ mutation eliminates the exon 6d containing isoforms only, C, F, G and H. Their depletion, however, is not enough to cause lethality. Concerning the genomic duplications, a similar graded array of normal functions can be identified. The large Dp3 is the most effective supplier of all normal functions, followed, in descending order, by Dp1 and Dp2. Although the three duplications can rescue all DL and *wupA* mutants in general, Dp2 does not cover *hdp*^*3*^ nor *hdp*^*2*^ alleles. The graded scale for *wupA* is equivalent to that observed for *dpp* alleles [45] (see **Suppl. Fig. S1**).

**Fig 8.**
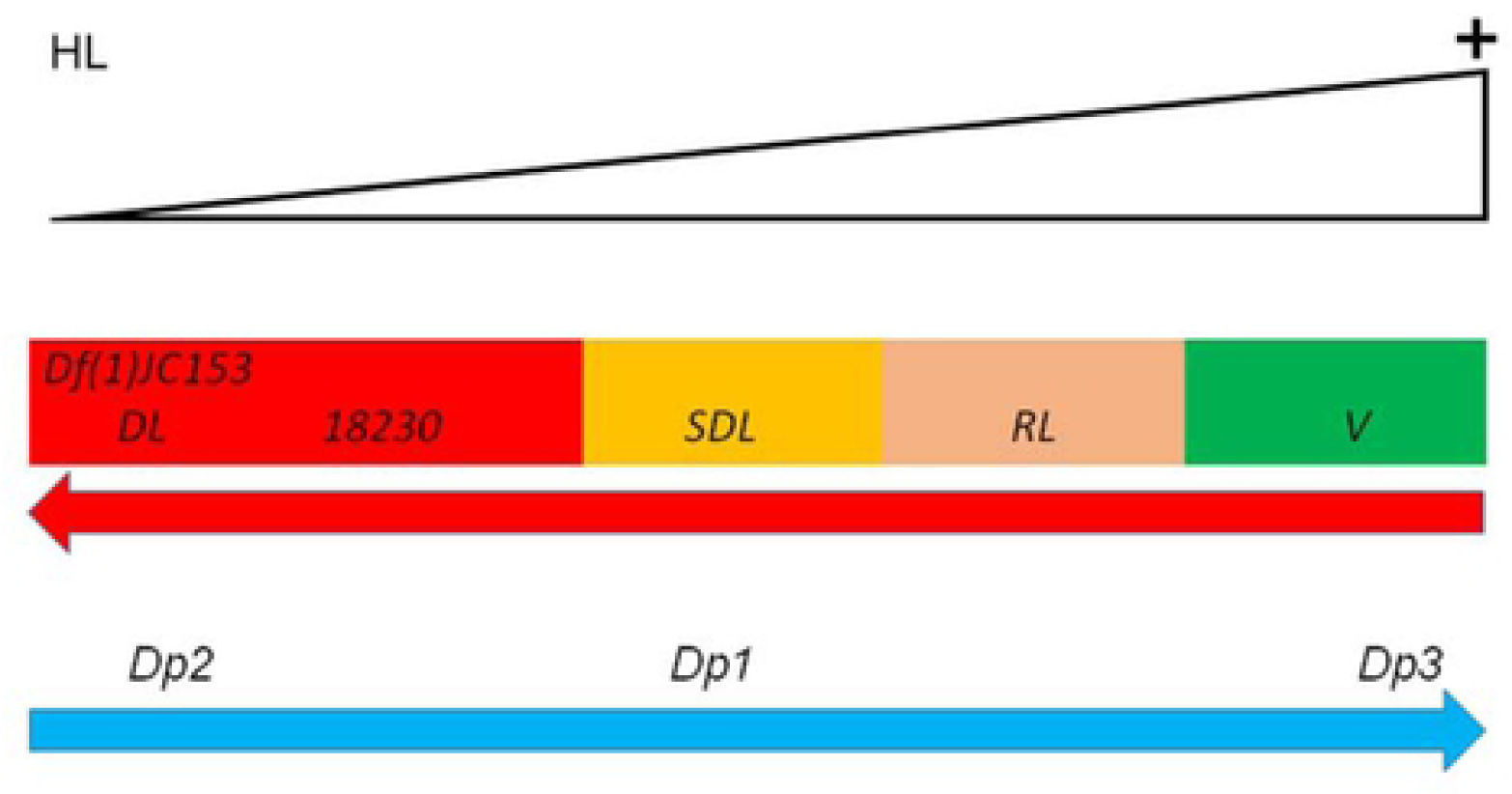
Graded scale of *wupA* mutant alleles. Schematic representation of the relative levels of *wupA* function in the corresponding mutant alleles and duplications.

In conclusion, the *wupA* gene encodes a number of TnI protein isoforms which show functional diversity and interactions. Their transcriptional expression is quantitatively regulated to the extent that their combined haploinsuficiency, at least all the red isoforms, causes haplolethality. By contrast, in the case of *dpp*, the haplolethality seems to result from the insufficiency of only one encoded protein, although five different transcripts mediate this expression [46]. The *wupA* and *dpp* associated HL functions seem functionally independent, among them and with respect to the other HL regions of *Drosophila*.

## DISCUSSION

Two major findings are reported here. A) TnI isoforms are functionally diverse and interacting, and B) This repertoire is strongly dosage sensitive for organism viability to the extent of causing haplolethality.

### Biology of TnI isoforms

TnI was once considered a muscle specific protein playing its role within the Troponin-Tropomyosin complex during contraction of the sarcomere [47]. Our previous studies demonstrated that TnI is also expressed in non-muscle cells and that, at least some protein isoforms, play a role in normal and tumorous cell proliferation and in apico-basal cell polarity [19,48]. Here, we analyzed the TnI isoforms separately, and found that their role in cell proliferation is diverse and, in some cases, opposed. This implies a wide network of functional interactions between TnI isoforms themselves and with other partners. When the coding sequence of *wupA* was identified [15], the array of alternative and mutually exclusive set of exons 6 a-d pointed to the possibility of functional specialization of TnI isoforms. One of the structural differences that the four alternative exons 6 offer is the number of Cys residues in each isoform. Differences in this number could provide different interactions with other Cys containing partners. This possibility should be explored in future protein-protein interaction studies using single TnI isoforms as bait.

We had shown previously that, at least some, TnI isoforms can translocate to the cell nucleus [18] where they elicit transcriptional changes in cell proliferation related genes [19]. Here we show that isoform K, and presumably the sequence identical L and M isoforms, can translocate to the nucleus. The high levels of Actin in the cell nucleus and its role in chromosome motion and gene transcription [49–51] is consistent with the proposal that the transcriptional changes elicited by the nuclear TnI could be mediated by the interaction of Actin with one or several TnI isoforms, akin to the interactions that take place in the muscle sarcomere. In the context of this study, at least isoforms A and B interfere with the nuclear trafficking of K. This feature provides a putative mechanism to influence cell proliferation through the quantitative regulation of the ratio between red and blue TnI isoforms according to the cell status or physiological requirements. Experiments to address this hypothesis should be designed under conditions that ensure the quantitative control of combinations of TnI isoforms. Such experiments, however, are beyond the scope of this study given the vast number of possible combinations. The functional diversity and interference among TnI isoforms implies the existence of regulatory mechanisms for their proper expression in the normal biology of the cell.

### Regulation of *wupA* expression

Previous to this study, we had identified two regulatory regions, URE and IRE, located at the 5’ terminus of *wupA*, based on the criteria of *LacZ*-reported expression of selected genomic fragments [17]. That criteria, however, is informative on positive enhancers, but repressors are missed. Here, we used qRT-PCR to discriminate among the various gene transcripts. The two genomic duplications analyzed, Dp1 and Dp2, although they rescue the DL mutants, the quantitative levels of transcripts they supply is heterogeneous with respect to the normal condition. This feature illustrates how superficial the evaluation of gene activity can be, if it is solely based on the resulting phenotype, adult viability in this case. Although these genomic duplications can restore viability of DL mutants, their transcription is far from being wild type. It is evident that the quantitatively normal levels of transcription require the proper chromatin landscape to an extent beyond the limits usually defined by a transcription unit. Additional supporting evidences can be found scattered in the scientific literature on other genes and organisms. However, these facts are largely unattended when experimenting with genomic fragments or, exceedingly so, when handling genetic constructs engineered with alien promoters and enhancers. The fact that none of the single isoforms rescue any of the known *wupA* mutants strongly suggests that the corresponding normal functions are achieved by combinations, rather than individual, isoforms acting in a specific stoichiometry. This is conceptually relevant because it underscores the combinatorial role of isoforms from a single gene, a subject often not considered in most studies.

The case of *hdp*^*3*^ has unveiled another unexpected feature on quantitative control of *wupA* transcription. A single nucleotide mutation at the splicing acceptor site for exon 6d increases the expression of non-exon 6d containing transcripts. These results on transcriptional activity indicate that the expression of *wupA* could be subject to a feed-back control by its own encoded products, in addition to the two ATG sites, and the duplicated regulatory 5’ regions, URE and IRE. The complexity of this regulation invites to re-consider the “regulatory hypothesis” for the HL function.

### The regulatory vs the functional hypotheses for the 16F7 HL function

Once the nature of the HL function at 16F7 is ascribed to the *wupA* gene exclusively, two possible mechanisms can be envisioned: A) A regulatory structure in the DNA located in the site affected by all DL rearrangement mutants, the interval between coordinates 1086.8 and 1086.9 towards the 3’ end of the gene, and B) Critical haploinsufficiency of several of the TnI isoforms. The cluster of DL rearrangements points to a 4Kb region in the intron between exons 7 and 8. Of note, the rearrangements leading to DL mutations have occurred in non-coding sequences in all HL associated genes known to date in flies and mice. This correlation could suggest that a certain regulatory structure in non-coding DNA might be responsible for the HL function through the proper transcriptional regulation of the products encoded in the corresponding gene.

This putative regulatory structure in *wupA* cannot operate in *trans* since the combination of the genomic fragments *E6L* plus *Dp*(*1;2R*) *CH322-143G12r* does not rescue DL mutants. Also, the URE and IRE regulatory regions cannot be the sole mechanism that regulates *wupA* expression because their structural alteration by means of deletions (*l*(*1*)*23437*) or insertions (*PL87* and *PG31*), yield RL, but not DL, phenotypes. The URE-IRE regions, however, exhibit chromosome pairing effects whereby the homozygosis for *PL87* restores normal expression of the gene [17]. Although the *LacZ* reported expression domains instructed by URE and IRE seem identical, they are not functionally redundant because the homozygosis for *l*(*1*)*23437*, which deletes only one domain, is still lethal [17]. Likewise, the homozygosis for any of the DL rearrangements maintains the dominant lethality. The clustering of DL rearrangements towards the 3’ end of *wupA*, plus the failure of incomplete genomic fragments to rescue DL phenotypes, suggests that the integrity of the transcription unit and, likely, some adjacent sequences, is required for the normal expression of the gene. This clustering leading to DL phenotypes would be akin to regulatory landscapes proposed for developmental genes [52]. Consistent with this proposal, the available database information on Hi-C domains [53,54] indicate that the *wupA* gene is contained within a single tridimensional chromatin domain (http://epigenomegateway.wustl.edu/browser). Thus, the native chromatin in and around *wupA* could be required to support the structural regulatory component of its HL function. The role of chromatin structure on the transcriptional regulation of adjacent genes, however, is still a matter of debate [55].

On the alternative hypothesis, the haploinsufficiency of one key TnI isoform, the individual overexpression of single TnI isoforms does not rescue the DL mutants. Thus, the HL function is clearly not the result of the depletion of just one isoform. By contrast, deleting the complete red subset of isoforms, as in the *18230* alleles, does yield a DL phenotype. This is the strongest evidence to support the functional hypothesis for this HL function. Nevertheless, could both hypotheses be reconciled? We reason that the proper regulation of *wupA* expression could be achieved by a structural organization that requires the integrity of the whole genomic region, in particular the intron between exons 7 and 8. This would explain why all DL mutants isolated so far are rearrangements affecting that intron. The mutants *18230C* and *18230B*, in effect, carry two mutations, the ATG>TGA and the insertion of the GFP reporter cassette in intron 1 in reverse orientation and driven by its own hsp60 promoter. For the reasons described above (**Fig. 8**), these mutants, although DL, they are less severe than the rearrangements clustered on the 7-8 intron. This is another evidence of the graded nature of haploinsufficiency, finally leading to dominant lethality.

Considering all available evidences, we interpret the HL function as a regulatory mechanism that ensures the proper expression of a gene, not only in time and space, but most significantly, in the required stoichiometry of the products encoded in the corresponding gene, *wupA* in the case studied here.

### Significance of HL functions

The location of *wupA* on the sexually dimorphic X chromosome may invite to consider the possibility that the HL condition could result from a special form of dosage compensation mechanism akin to those already known to tune up the transcription of X-linked genes in males [56–59]. However, *Drosophila* HL regions can be found in the X and autosomes. Also, our previous experiments to address the possibility of altered dosage compensation for *wupA* mutants yielded negative results [12].

Quantitative regulation of gene transcripts can be achieved by a number of mechanisms, including their codon identity composition, which affects their stability [60], or through modified tRNAs, which affect their translational efficiency [61]. In *wupA*, however, the normal HL function depends on a combination of, rather than single, transcripts. Genomes differ in their DNA content, both between cells and between individuals, and this variation is thought to contribute to adaptation and evolution [62–65]. However, DL mutants are not an extreme case of dominant mutations nor have the possibility of being positively selected. From yeast to flies, the haploid condition for genes encoding proteins with very general functions in cell biology shows a phenotype of haploinsufficiency. For instance, *Minute* genes in *Drosophila* usually encode proteins involved in translation or general metabolism, and their haploinsufficiency causes developmental delay and reduced size, but not dominant lethality [63]. Equivalent to fly *Minutes*, the 170 haploinsufficient genes in yeast can be recovered as viable organisms if the culture medium is metabolically adjusted [5].

What do HL functions have in common? The HL function of *wupA*, and to some extent that of *dpp*, are the only cases for which a mutational analysis has been carried out. Two features are in common in both cases, the corresponding DL mutants are chromosomal rearrangements, and the affected gene encodes several transcripts. However, functional diversity of products from HL associated genes has been studied for *wupA* only. It could be argued that one or several gene products are required in a critical amount for viability. The case of *wupA* HL shows (**Table 2**) that none of the transcriptional products, taken one at a time, can rescue the DL mutants; thus, it is clear that the hypothetically critical amount for viability should correspond to more than one product. We co-overexpressed several of the UAS constructs (see foot note on **Table 2**) but none of them rescued the DL phenotype. Only when all red subset of *wupA* products are eliminated (mutants *wupA18320C* and *wupA18320B*), a DL phenotype is obtained.

The fly genes *wupA* and *dpp* have homologues in humans, *TNNI* and *BMP*. While *wupA* encodes 11 TnI protein isoforms, the TNNI human counterparts are encoded in three different genes, *TNNI1, TNNI2* and *TNNI3*. Based on the association between copy number variation and certain types of cancer (Catalogue of Somatic Mutations in Cancer: https://cancer.sanger.ac.uk/cosmic), and experimental tumor cell growth suppression, *TNNI1* seems the closest homologue to *wupA* [18]. The sequence similarity between the three human genes versus the single fly gen, however, is not very different (31, 32 and 34%, respectively). The corresponding chromosomal locations, 1q32.1, 11p15.5 and 19q13.42, respectively, are not included in the regions never recovered in haploid condition. Nevertheless, whole genome analysis of the predicted probability of being haploinsufficient, indicates that *TNNI3* has a high probability [1]. A protein interactome data base (http://www.interactome-atlas.org/search) shows that the three human TNNI interact with several proteins beyond the classical components of the muscle sarcomere. Notably, TNNI1 exhibits the largest repertoire of interactions including, among others, RPAC1, a DNA-dependent RNA polymerase, and CDC7AL, a cell cycle associated protein. These features are consistent with the transcriptional and cell proliferation effects identified here for *Drosophila* TnI.

As for dpp/BMP, although the fly gene encodes 1 protein through 5 transcripts, the human homologue is represented by a family of 10 members, from which *BMP2* and *BMP4* can be considered the closest to *dpp*. The murine and human homologs of *Vegf* and *Tcof1* have in common the coding for several proteins, and the presence of multiple (*Vegf*) or two (*Tcof1*) ATG sites (www.Ensembl.org). They have a relatively high probability of being haploinsufficient [1], albeit the probability of being haplolethal cannot be estimated currently.

One would have expected that HL functions will be under such a negative selective pressure as to have been eliminated. Yet, they seem to exist in far apart species; *Drosophila* contains five of them, and mice at least two. Actually, this paradox extends to haploinsufficient genes since they represent a barrier to organismal fitness due to the resulting pathologies. Haploinsufficiency cannot be overcome by compensatory overexpression of related genes because these transcriptional network effects are rare [31]. A dosage-stabilizing hypothesis has been proposed to explain the persistence of haploinsufficient genes, based on the possibility that, both in their attenuated and excess of function conditions are deleterious for the organism [64]. Although we do not argue against this proposal, we find it unlikely for haplolethality because large duplications covering each of these fly HL regions, except Tpl, are viable. We speculate that HL functions could be maintained as a result of a complex regulatory mechanism of genes whose encoded products need to be expressed in specific quantities and combinations because of their functional interactions in normal biology. Only when the repertoire of products splits into separate genes, as after gene duplication, the initial HL function would dissociate from the initial single gene. That dissociation, however, would be apparent only. Likely, the combined haploidy of the duplicated genes would reveal again the original haplolethality. Under this hypothesis, haplolethality would not be a property of the gene, but a property of the quantitative requirements of the encoded products. A systematic search for HL functions in a particular genome could unveil regulatory interactions unsuspected hitherto. In all likelihood, the HL condition of *wupA* in *Drosophila* is not unique across genomes. However, to find out if other HL regions result from a quantitative regulation of combinations of gene products, equivalent studies to those carried out here will be necessary.

## MATERIALS AND METHODS

### Mutant strains

Flies were raised in standard fly food at 25°C. As *wupA* mutations we used the following fly stocks from our own collection: *Df*(*1*) *TnI*^*23437*^, *In*(*1*)*PL87* and *In*(*1*)*PG31* [19] are three rearrangements located the 5’ URE and IRE regulatory region of the gene [17] (see also **Fig. 2**). Other mutant alleles and genomic constructs have been described previously [12,16]. To elicit excess of *wupA* function, we used the *PBac*(*WH*)*f06492* construct from Exelixis referred here as *UAS-TnI*^*f06492*^. The genomic fragments *CH322-143G12* (22251 bp), *CH322-61E02r* (20843 bp) or *CH322-04P16r* (18542 bp) were generated in [67]. The three fragments were cloned in the vector attB-P[acman]-CmR-BW-F-2-attB-BW3 (accession FJ931533). Each bacterial artificial chromosome (BAC) was injected in embryos of the stock *y*^1^ *M*{*vas-int*.*Dm*}*ZH-2A w*[*]; *M*{*3xP3-RFP*.*attP’*}*ZH-22A* (BL24481) and adult transgenic flies were selected by the reporter RFP signal in the oceli. To analyze other HL regions, the following stocks were obtained from the Bloomington collection: *C*(*X;Y*), *y sn Grip91/C*(*1*)*RM, y v*; *Dp*(*1;f*)*LJ9, y*^+^ (BL5128); *Dp*(*2;1*)*G146,dpp*^+^*/FM7i*; *dpp*^*H46*^ *wg*^*Sp*^ *cn bw/CyO* (BL2060); *Df*(*1*)*hl-a, w cv B/FM6*; *Dp*(*1;2*)*sn*^+*72d*^/+ (BL6698) and *Dp*(*1;2*)*CH322-143G12r/CyO*; *UAS*(*y*^+^*v*^+^) *up*^*RNAi*^*attP2/TM6* (from BL31541).

### Generation of UAS-*wupA* transgenic lines

All DNA plasmids were generated by RECOMBINA S.L. (Navarra, Spain). Full length DNA sequence of each *wupA* transcript was amplified by PCR. The products were cloned in pENTRY vector as an intermediate step. Then, each *wupA* fragment was subcloned in pUASp vector via NotI/XbaI restriction enzymes and injected in *y w* embryos. For the HA-tagged version of the K isoform, the *wupA-K* coding sequence was amplified by PCR. The primers *wupA-K-HA* forward and *wupA-K-HA* reverse introduced an in-frame HA epitope coding sequence in 3’. The fragments were cloned in pENTRY vector and subcloned in pUASp vector via KpnI/XbaI restriction enzymes and injected in *y w* embryos. Sequence primers were as follows: *wupA-K-HA* F>cggGGTACCatggaggaagcctccaaggccaa *wupA-K-HA R*>tagTCTAGAttaAGCGTAATCTGGTACGTCGTATGGGTAagcttcggcctcaacctcct

### Somatic clone induction and quantification

Crosses were set with 10 females and 10 males per vial at 25°C changed every 72 hours to avoid overcrowding. FLP-out clones were obtained by delivering a heat shock (8 minutes at 37°C) during 2^nd^ instar larvae of *HsFlp; Act-FRT-yellow-FRT-Gal4;* + crossed against the corresponding UAS-*wupA* transgenic line isoform (A, - B, -C, -D, -E, -F, -G, -H, -I, -J or -K). 48 hours after heat shock, 3^rd^ instar wandering larvae were dissected for clone screening. Control cultures (UAS-*LacZ*) were run in parallel. A software-assisted area measurement (Bitplane’s Imaris Surface) was used to obtain clone area and cell size. Cell profiles are identified by the myrRFP reporter and cell nuclei are revealed by DAPI. The “surface” option was chosen to measure the areas occupied by DAPI or RFP pixels from the entire 3D image. The area of the clone (RFP-marked) was calculated and divided by DAPI-marked area. As a result, the percentage of wing disc occupied by genetically marked cells was represented.

### Viability quantification

To determine the rescue of dominant lethal phenotypes, we performed viability assays. 10 females and 10 males were crossed and maintained in the same tube for 72 hours, then adult flies were changed to a new tube and embryos were incubated at 25°C. The total number of adult flies was counted in three independent experiments. The number of experimental adult flies was divided by the number of control siblings (balancer) adult flies to indicate the survival ratio.

### Quantitative PCR assays

For qRT-PCR assays, RNA was extracted with Trizol (Invitrogen) according to standard procedures. To prevent genomic DNA contamination all RNA samples were treated with DNaseI according to manufacturer’s procedures (30 min at 37°C). Primers were designed to anneal in different exons from each gene, later qPCR products were run in 2% agarose gels to analyze the bands. All the products correspond with their expected length, which indicates that no genomic DNA contamination was present. Assays were performed in triplicates using RNApolII as a housekeeping gene. Drosophila Troponin-I TaqMan Gene Expression probe (Applied Biosystems) was used. Used primers were:

wupA 5-6F GGCTAAACAGGCTGAGATCG

wupA 6a R TCGATGATGCGTCTACGTTC

wupA 6b R TCAACATCCTTGCGTTTAACA

wupA 6c R AAATCGTACTTTTCGGACTCCA

wupA 6d R AGATCCCATTTCTGGCCTTC

wupA 11/12 F GCCCAAGTTAACGATCTTCG

wupA 11/12 R TCCAGCGTGAACTCCTTCTT

RpL32 F TGTCCTTCCAGCTTCAAGAT

RpL32 R CTTGGGCTTGCGCCATTTG

### Statistics

Statistical significance was calculated with the two-tailed Student’s *t*-test or ANOVA test. Significance levels are indicated as *p < 0,05; **p < 0,005 or ***p < 0,001. Number of samples N > 8 animals in confocal imaging, and n=3 sets of 5 adults, male or female as indicated, for qPCR experiments.

## ACKNOWLEDGMENTS

Research was funded by grants from the Spanish Ministry of Economy (PGC2018-094630-B-100, BFU2015-65685-P and PID2019-110116GB-100). We are grateful to the Bloomington Drosophila Stock Center (NIH P40OD018537) for fly strains. Transgenic lines were generated by RECOMBINA S.L. (Navarra, Spain) and the CRISPR alleles *18320B* and *18320C* were produced by WellGenetics Inc. (Taiwan). Ania Angüi provided the data for Fig. 5A. The technical help of Esther Seco and Daniela Escobar and critical comments from lab members are most appreciated.

## SUPPLEMENTARY DATA

### Supplementary figure legends

**Fig. S1.- Location of dominant lethal rearrangement in the *dpp* gene**. Data taken from Fig 6 of [11]. The location of the small deletion causing the *dpp*^*H46*^ allele is described in [46]. The graded allelic series of *dpp* alleles is reported in [45]. It should be noted that that report describes 5 transcripts emerging from the *dpp* gene although the current data base of FlyBase indicates only four.

**Fig. S2**.- Set of genes tested in the qRT-PCR assays of *tub-Gal4*^*LL7*^>*UAS-TnI-K* female larvae using *tub-Gal4*^*LL7*^>*UAS-LacZ* female larvae from a parallel cross as control. All assay determinations were done in triplicate.

**Fig. S3.- Overexpression of TnI isoform K in wing domains cause morphological abnormalities**. Adult wings expressing isoform K in the *rn-Gal4* domain. Note the curved wings (arrows). These adults exhibited low viability with respect to siblings.

**Fig. S4.- Design of CRISPR/Cas9 mutations in the red ATG site of *wupA***. Mutations *18320B* and *18320C* were produced by WellGenetics Inc. (Taiwan). The later was validated by sequencing.

